# Mucosal antibody responses to SARS-CoV-2 booster vaccination and breakthrough infection

**DOI:** 10.1101/2023.08.24.554732

**Authors:** Disha Bhavsar, Gagandeep Singh, Kaori Sano, Charles Gleason, Komal Srivastava, PARIS Study Group, Juan Manuel Carreño, Viviana Simon, Florian Krammer

## Abstract

Coronavirus disease 2019 (COVID-19) vaccines have saved millions of lives. However, variants of severe acute respiratory syndrome coronavirus 2 (SARS-CoV-2) have emerged causing large numbers of breakthrough infections. These developments necessitated the rollout of COVID-19 vaccine booster doses. It has been reported that mucosal antibody levels in the upper respiratory tract, especially for secretory IgA (sIgA), correlate with protection from infection with SARS-CoV-2. However, it is still unclear how high levels of mucosal antibodies can be induced. In this study, we measured serum IgG, saliva IgG and saliva sIgA responses in individuals who received COVID-19 mRNA booster vaccinations or who experienced breakthrough infections. We found that mRNA booster doses could induce robust serum and saliva IgG responses, especially in individuals who had not experienced infections before, but saliva sIgA responses were weak. In contrast, breakthrough infections in individuals who had received the primary mRNA vaccination series induced robust serum and saliva IgG as well as saliva sIgA responses. Individuals who had received a booster dose and then had a breakthrough infection showed low IgG induction in serum and saliva but still responded with robust saliva sIgA induction. These data suggest that upper respiratory tract exposure to antigen is an efficient way of inducing mucosal sIgA while exposure via intramuscular injection is not.

**Importance:** Antibodies on mucosal surfaces of the upper respiratory tract have been shown to be important for protection from infection with SARS-CoV-2. Here we investigate the induction of serum IgG, saliva IgG and saliva sIgA after COVID-19 mRNA booster vaccination or breakthrough infections.

## Observation

Vaccines for coronavirus disease 2019 (COVID-19) have saved millions of lives (1). However, variants of severe acute respiratory syndrome coronavirus 2 (SARS-CoV-2) that escape neutralizing antibody responses have significantly decreased vaccine effectiveness against both infection and symptomatic disease (2). Importantly, in order to protect from infection, robust titers of neutralizing mucosal antibodies in the upper respiratory tract are likely needed. In fact, mucosal antibody titers have been linked to protection from infection (3-5). However, injected vaccines are not very effective in inducing mucosal immunity in the upper respiratory tract (6). Right after vaccination, when IgG serum titers are very high, IgG titers sufficient for protection are likely present on mucosal surfaces of the upper respiratory tract too. However, as titers wane, these IgG levels likely decline to sub-protective levels. Initial efficacy of protection from infection of the mRNA vaccines which declined quickly support this hypothesis (7). However, secretory IgA (sIgA), which is actively transported to mucosal surfaces and is produced by B cells in the *lamina propria* below these surfaces are typically not induced by vaccination. It is believed that these mucosa-specific sIgA responses are mostly induced when antigen is encountered via the mucosal route, e.g. after infection with a respiratory virus. We have in the past shown that mRNA vaccination in previously naïve individuals results in mucosal IgG but not mucosal sIgA responses while sIgA was induced by mRNA vaccination in individuals who already had been infected before vaccination with SARS-CoV-2 (6). Here we explored the induction of mucosal IgG and sIgA responses in individuals who received booster vaccine doses (3^rd^ dose) or who had breakthrough infections after the primary vaccination series in our PARIS (Protection Associated with Rapid Immunity to SARS-CoV-2) cohort (8).

### Mucosal antibody responses after mRNA booster dose

Our cohort included individuals who were naïve before they received the primary COVID-19 mRNA vaccination series (n=30, 18 females) and individuals who were infected and then received the primary COVID-19 mRNA vaccination series (hybrid immune; n=30, 19 females). We wanted to see what happened to serum and mucosal antibody titers when these individuals received the booster dose (3^rd^ dose) of mRNA vaccine. For previously naïve individuals pre-boost samples were taken on a median 8 days before vaccination (range 0-30 days) and post-boost samples were taken on a median 27 days after vaccination (range 20-42 days). For hybrid immune individuals samples were taken on a median 11.5 days (range 0-50 days) before the booster dose and on a median 28 days (range 18-57 days) after the booster dose. Individuals without prior infection responded with a robust serum IgG response (**Figure 1A** and **D**, 30.2-fold induction) and saliva IgG titer increase as well (**Figure 1B** and **D**, 10.5-fold). While mucosal sIgA were also significantly induced (**Figure 1C** and **D**), this induction was much less robust and only 2.5-fold. For hybrid immune individuals, we found a lower level of induction of IgG in serum (**Figure 1E** and **H**, 4.0-fold) and saliva (**Figure 1F** and H, 2.9-fold), likely due to higher pre-boost titers. Saliva sIgA was induced but only 2.1-fold (**Figure 1G** and **H**).

**Figure 1.**
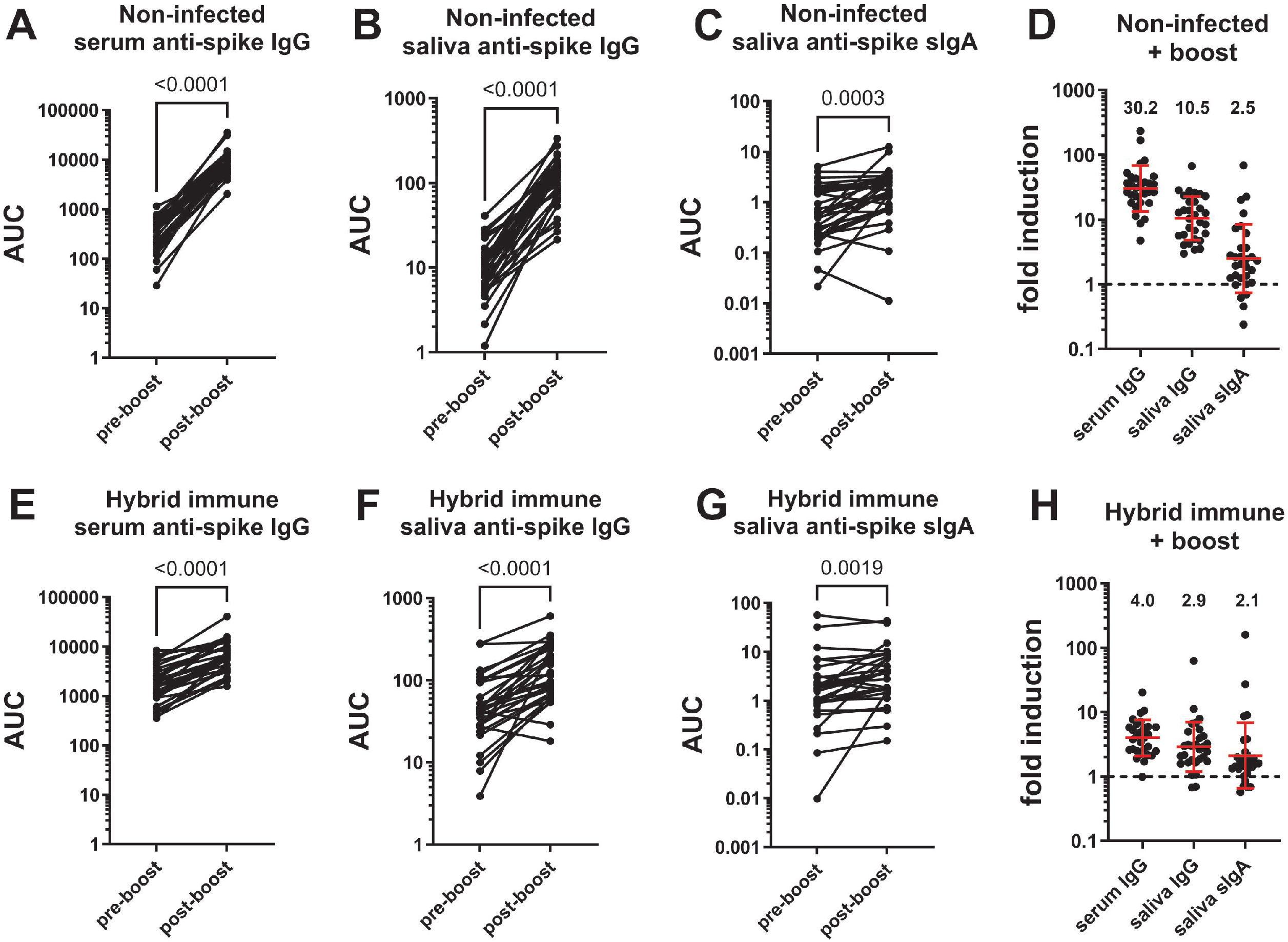
Induction of serum IgG, saliva IgG and saliva sIgA after COVID-19 mRNA booster vaccination. Pre- and post-boost serum IgG (**A**), saliva IgG (**B**) and saliva sIgA (**C**) titers in non-infected individuals who received the primary vaccination series. (**D**) shows fold induction of absolute titers presented in A, B and C. Pre- and post-boost serum IgG (**E**), saliva IgG (**F**) and saliva sIgA (**G**) titers in hybrid immune individuals who received the primary vaccination series. (**H**) shows fold induction of absolute titers presented in A, B and C. AUC= area under the curve. Statistical analysis in A, B, C, E, F and G was performed using a ratio paired t test. The red bar in D and H indicates the geometric mean, the error bars indicate the standard deviation of the geometric mean. The dotted lines indicate no induction (1-fold). N=29-30 for each panel.

### Mucosal antibody responses after SARS-CoV-2 breakthrough infections

As a next step, we wanted to evaluate the serum IgG, saliva IgG and saliva sIgA response after breakthrough infections. Here, we split out participants into individuals who had their breakthrough infections after the primary mRNA COVID-19 vaccination series (n=17, 13 females, 3 Delta breakthroughs, 14 Omicron breakthroughs) or after the mRNA COVID-19 booster dose (n=34, 26 females, all Omicron breakthroughs). For individuals vaccinated only with the primary vaccination series pre-breakthrough samples were taken on a median 282 days before vaccination (range 37-341 days) and post-breakthrough samples were taken on a median 30 days after vaccination (range 8-59 days). For boosted individuals samples were taken on a median 55.5 days (range 8-129 days) before the booster dose and a median 29.5 days (range 12-48 days) after the booster dose. For breakthrough infections after the primary vaccination series we found robust induction of serum IgG, saliva IgG and saliva sIgA (**Figure 2A-D**) with 10.6-fold, 11.9-fold and 11.1-fold induction, respectively. Antibody induction in breakthrough cases after the booster dose was detectible for serum IgG and saliva IgG but less pronounced with 2.0-fold and 1.8-fold inductions respectively (**Figure 2E, F** and **H**). However, the sIgA induction was still robust (6.9-fold) but was very heterogeneous (**Figure 2G** and **H**).

**Figure 2.**
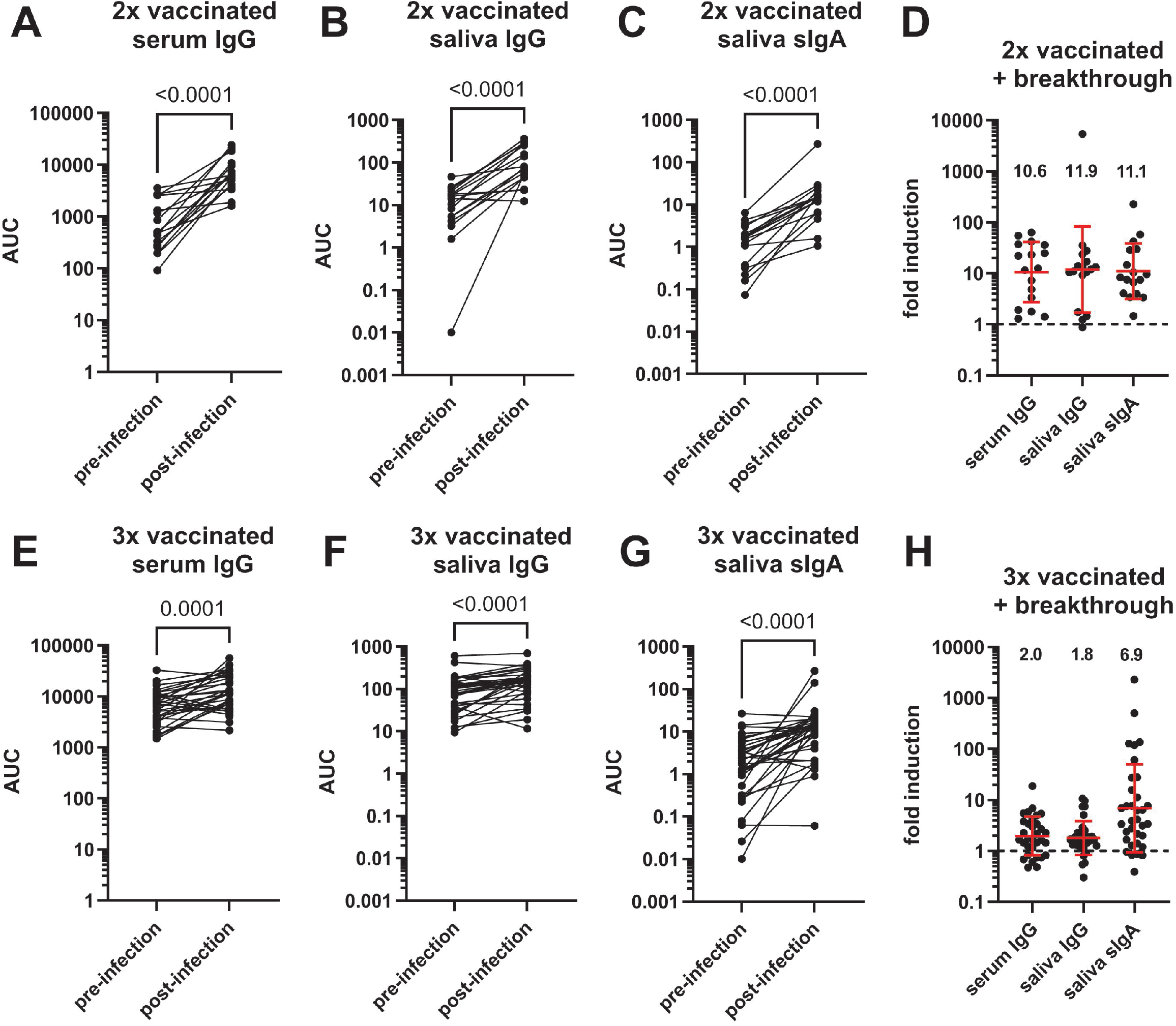
Induction of serum IgG, saliva IgG and saliva sIgA after SARS-CoV-2 breakthrough infections. Pre- and post-breakthrough serum IgG (**A**), saliva IgG (**B**) and saliva sIgA (**C**) titers in individuals who received the primary vaccination series. (**D**) shows fold induction of absolute titers presented in A, B and C. Pre- and post-breakthrough serum IgG (**E**), saliva IgG (**F**) and saliva sIgA (**G**) titers in individuals who had their breakthrough infections after the booster dose. (**H**) shows fold induction of absolute titers presented in A, B and C. AUC= area under the curve. Statistical analysis in A, B, C, E, F and G was performed using a ratio paired t test. The red bar in D and H indicates the geometric mean, the error bars indicate the standard deviation of the geometric mean. The dotted lines indicate no induction (1-fold). N=17 for panels A, B, C and D and N=32-34 for the remaining panels.

## Conclusions

In summary, we observed that booster vaccination in previously naïve individuals induced strong serum IgG responses and saliva IgG responses but the induction of saliva sIgA was low. In individuals with hybrid immunity, antibody induction after the booster dose was lower in general likely owing to higher baseline titers. Breakthrough infections after the primary vaccination series resulted in robust induction of serum IgG, saliva IgG and saliva sIgA. However, breakthrough infection after the booster dose led to a lower level of IgG induction in serum and saliva while sIgA induction in saliva was still robust, even though there was a lot of variation among individuals. Our data suggest that breakthrough infections induce robust mucosal sIgA while injected booster doses of COVID-19 mRNA vaccines do not. This is in line with the concept that mucosal antigen delivery is needed for efficient induction of sIgA in the upper respiratory tract.

## Acknowledgments

We thank all the participants of our longitudinal PARIS study for their generous and continued support of research. This effort was supported by the Serological Sciences Network (SeroNet) in part with Federal funds from the National Cancer Institute, National Institutes of Health, under Contract No. 75N91019D00024, Task Order No. 75N91021F00001. The content of this publication does not necessarily reflect the views or policies of the Department of Health and Human Services, nor does mention of trade names, commercial products or organizations imply endorsement by the U.S. Government. This work was also partially funded by the Centers of Excellence for Influenza Research and Surveillance (CEIRS, contract # HHSN272201400008C), the Centers of Excellence for Influenza Research and Response (CEIRR, contract # 75N93021C00014), by the Collaborative Influenza Vaccine Innovation Centers (CIVICs contract # 75N93019C00051) and by institutional funds. KS was supported by the Japanese Society for the Promotion of Science (JSPS) Overseas Research Fellowship.

## Data availability statement

All data will be available from ImmPort under the following identifiers: XXX.

## Supplemental Material

### Methods and materials

#### Human samples (Participant enrollment, serum and saliva collection and methods)

Serum and saliva samples were collected from participants in the longitudinal observational PARIS (Protection Associated with Rapid Immunity to SARS-CoV-2) study (8). The study was reviewed and approved by the Mount Sinai Hospital Institutional Review Board (IRB-20-03374/Study-20-00442). All participants signed written consent forms prior to sample and data collection. All participants provided permission for sample banking and sharing.

#### Recombinant proteins

Recombinant SARS-CoV-2 Spike (S) proteins were produced using a mammalian cell protein expression system as described in detail by (9, 10). In brief, the S gene sequence of the ancestral spike of SARS-CoV-2 (GenBank: MN908947) was modified to include six prolines (HexaPro (11)) and cloned into the mammalian expression vector pCAGGs. The spike was then expressed using the Expi293 Expression System (Thermo Fisher Scientific), according to the manufacturer’s instructions. Cell culture supernatants were centrifuged at 4000 x g, filtered, and purified with Ni-nitriloacetic acid (NTA) agarose (Qiagen). Purified S protein was concentrated and buffer was exchanged to phosphate buffered saline (PBS) (pH7.4) using Amicon Ultracell (Merck) centrifugation units. Proteins were stored at -80°C until use.

#### Antigen specific IgG and sIgA ELISA

Immulon 4 HBX 96-well microtiter plates (Thermo Fisher Scientific, #3855) were coated with recombinant proteins (100 ng/well) in PBS (pH 7.4) overnight at 4°C. Well contents were discarded and blocked with 200 μl of 3% non-fat milk (AmericanBio, # AB10109-01000,) in PBS containing 0.1% Tween-20 (PBST) (5% non-fat milk in PBST for saliva IgG and sIgA) for one hour at room temperature (RT). After blocking, 100 μl of serum samples (starting at 1:80 and serially diluted three-fold with 1% non-fat milk in PBST) or 50 μl of saliva samples (starting at 1:2 and serially diluted two-fold with 2.5% non-fat milk in PBST) were added to each well. Samples were incubated either at RT for two hours or overnight at 4°C for IgG or sIgA measurement and plates were washed with PBST three times. For serum IgG measurement, 50 μl of HRP-labeled goat anti-human IgG antibody (Sigma-Aldrich, #A0170) diluted 3000-fold with 1% non-fat milk in PBST were added to each well and incubated at RT for 1 hour. For saliva IgG measurement, 50 μl of HRP-labeled goat anti-human IgG antibody (Invitrogen, #31412) diluted to 0.32 mg/ml with 2.5% non-fat milk in PBST were added to each well and incubated at RT for 1 hour. For sIgA measurement, 50 μl of mouse anti-human secretory IgA antibody (MilliporeSigma, #HP6141) diluted to 5 μg/ml with 2.5% non-fat milk in PBST was added to each well and incubated at RT for 2 hours. These plates were washed again with PBST three times, and 50 μl of HRP labeled goat anti-mouse IgG Fc antibody (Invitrogen, #31439) diluted to 1:1000 with 2.5% non-fat milk in PBST was added to each well and incubated at RT for 1 hour. For all measurements, after final washing step with PBST three times, 100 μl of SIGMAFAST o-phenylenediamine dihydrochloride substrate solution (Sigma-Aldrich, #P9187) were added to each well at RT for 10 minutes. The reaction was stopped by addition of 50 μl of 3M hydrochloric acid (HCl). Optical density at 490 nm was measured using Synergy 4 (BioTek) plate reader. Eight wells on each plate received no primary antibody (blank wells) and the optical density in those wells was used to assess background. Area under the curve (AUC) was calculated by deducting the average of blank values plus three times standard deviation of the blank values. Antigen specific sIgA AUC values were adjusted by dividing the values by total IgA concentration within saliva samples (adjusted AUC).

#### Measurement of total IgA concentration within saliva samples

Immulon 4 HBX 96-well microtiter plates were coated with 250 ng/well of goat anti-human IgA (Bethyl Laboratories, #A80-102A) in PBS (pH 7.4) overnight at 4°C. Well contents were discarded and 200 μL/well of 5% non-fat milk in PBST were added and plates were incubated for one hour at RT. After blocking, 50 μL of saliva samples diluted (starting at 1:256 and serially diluted three-fold) in 2.5% non-fat milk PBST were added to each well and incubated at RT for two hours. Plates were washed with PBST three times and 50 μl/well of HRP-labeled goat anti-human IgA antibody (Bethyl Laboratories, #A80-102P) diluted to 1:10,000 in 2.5% non-fat milk in PBST was added to each well and incubated at RT for 1 hour. After a final washing step, 100 μl of SIGMAFAST o-phenylenediamine dihydrochloride substrate solution were added to each well for the reaction at RT for 10 minutes. Reaction was stopped by addition of 50 μl/well of 3M HCl. Optical density at 490 nm was measured using Synergy 4 plate reader. Purified human plasma IgA (EMD Millipore, #401098) was diluted (starting at 500 ng/ml and serially diluted two-fold) and was used as standard. A five-parameter logistic fit was conducted on the standard curve and IgA concentrations within saliva samples were calculated.

#### Statistical analysis

Differences between antibody titers between two groups was analyzed with a ratio paired t test. All statistical analyses, AUC calculation, and five-parameter logistic fit of standard curves were conducted with Graphpad Prism version 10.

